# SARS-CoV-2 RBD219-N1C1: A Yeast-Expressed SARS-CoV-2 Recombinant Receptor-Binding Domain Candidate Vaccine Stimulates Virus Neutralizing Antibodies and T-cell Immunity in Mice

**DOI:** 10.1101/2020.11.04.367359

**Authors:** Jeroen Pollet, Wen-Hsiang Chen, Leroy Versteeg, Brian Keegan, Bin Zhan, Junfei Wei, Zhuyun Liu, Jungsoon Lee, Rahki Kundu, Rakesh Adhikari, Cristina Poveda, Maria Jose Villar, Ana Carolina de Araujo Leao, Joanne Altieri Rivera, Zoha Momin, Portia M. Gillespie, Jason T. Kimata, Ulrich Strych, Peter J. Hotez, Maria Elena Bottazzi

## Abstract

There is an urgent need for an accessible and low-cost COVID-19 vaccine suitable for low- and middle-income countries. Here we report on the development of a SARS-CoV-2 receptor-binding domain (RBD) protein, expressed at high levels in yeast (*Pichia pastoris*), as a suitable vaccine candidate against COVID-19. After introducing two modifications into the wild-type RBD gene to reduce yeast-derived hyperglycosylation and improve stability during protein expression, we show that the recombinant protein, RBD219-N1C1, is equivalent to the wild-type RBD recombinant protein (RBD219-WT) in an *in vitro* ACE-2 binding assay. Immunogenicity studies of RBD219-N1C1 and RBD219-WT proteins formulated with Alhydrogel^®^ were conducted in mice, and, after two doses, both the RBD219-WT and RBD219-N1C1 vaccines induced high levels of binding IgG antibodies. Using a SARS-CoV-2 pseudovirus, we further showed that sera obtained after a two-dose immunization schedule of the vaccines were sufficient to elicit strong neutralizing antibody titers in the 1:1,000 to 1:10,000 range, for both antigens tested. The vaccines induced IFN-γ, IL-6, and IL-10 secretion, among other cytokines. Overall, these data suggest that the RBD219-N1C1 recombinant protein, produced in yeast, is suitable for further evaluation as a human COVID-19 vaccine, in particular, in an Alhydrogel^®^ containing formulation and possibly in combination with other immunostimulants.

## 1. Introduction

The number of coronavirus disease 19 (COVID-19) cases globally is readily over the 100-million-person mark, with over 2 million deaths up to early February 2021. In response to the pandemic, several vaccines have been shown to be effective and have been either approved or authorized for emergency use in different countries^1^. There are many ways to categorize the more than 100 potential COVID-19 vaccine candidates^2^, but one approach is to divide them as those employing novel, not hitherto licensed technologies for production, versus those employing traditional vaccine production approaches with precedence in licensed vaccines^3^. The Operation Warp Speed (OWS) initiative in the United States^4^ and similar efforts in other parts of the world^5^ initially focussed on approaches employing new platforms, including several messenger RNA (mRNA)-based vaccines as well as non-replicating adenovirus vector vaccines^4^. Among the more established or traditional approaches, whole-inactivated virus vaccines on aluminum oxy-hydroxide have been developed in China^6^, as have several recombinant protein vaccine candidates^7–10^. Each of these approaches offers both distinct advantages and disadvantages in terms of production, scale-up, potential efficacy, and safety, as well as delivery.

We have previously reported on recombinant protein-based coronavirus vaccine candidates, formulated with Alhydrogel^®^ to prevent Severe Acute Respiratory Syndrome (SARS)^11–13^ and Middle East Respiratory Syndrome (MERS)^14^. In both cases, the Receptor-Binding Domain (RBD) of the SARS or MERS spike proteins was used as the target vaccine antigen. In a mouse model, the SARS-CoV RBD219-N1/Alhydrogel^®^ vaccine induced high titers of virus-neutralizing antibodies and protective immunity against a mouse-adapted SARS-CoV virus challenge. It was also found to minimize or prevent eosinophilic immune enhancement when compared with the full spike protein^11^.

The RBD of SARS-CoV-2 has likewise attracted interest from several groups now entering clinical trials^8, 15–19^. Our original approach was to apply the lessons learned from SARS-CoV and accelerate COVID-19 vaccine development efforts using microbial fermentation in the yeast *Pichia pastoris*^*20*^. We selected the SARS-CoV-2 RBD219-WT sequence, residues 331-549, through alignment with the SARS-CoV RBD sequence, and later developed a genetically engineered version (RBD219-N1C1) to improve antigen stability^21^.

Yeast expression technology is widely available and used to produce, for example, the VLP antigen of the licensed HPV vaccine as well as recombinant hepatitis B vaccines in several middle-income countries (LMICs)^22^, including Bangladesh, Brazil, Cuba, India, and Indonesia over the last 35 years.

As COVID-19 spreads across the globe, especially among urban populations living in extreme poverty^23^, there will be greater urgency to produce safe, effective, highly scalable, and affordable COVID-19 vaccines locally or regionally. Therefore, the development of a yeast-expressed recombinant protein-based COVID-19 vaccine allows developing it for global health and populations vulnerable to poverty-related diseases^22^.

Here, we present the first preclinical data of a COVID-19 recombinant protein-based vaccine candidate, SARS-CoV2 RBD219-N1C1, formulated with Alhydrogel^®^. We demonstrate that modifications made to the SARS-CoV2 RBD gene to improve production and stability preserve the protein antigen functionality and its immunogenicity after Alhydrogel^®^ adsorption.

## 2. Materials and Methods

### 2.1 Cloning and expression of the genes encoding RBD219-WT and RBD219-N1C1

The RBD219-WT recombinant subunit protein contains amino acid residues 331-549 of the SARS-CoV-2 spike protein (GenBank No.: QHD43416.1). It contains a hexahistidine tag at its C-terminus. In the tag-free RBD219-N1C1 antigen candidate, the N331 glycosylation site has been removed, and C538 has been mutated to an alanine residue to prevent aggregation due to intermolecular disulfide bonding. The DNAs for both antigen candidates were individually synthesized with their codon use optimized for translation in *Pichia pastoris* and ligated into pPICZαA using the EcoRI and XbaI restriction sites (GenScript). The recombinant plasmids were electroporated into *P. pastoris* X33 following the EasySelect™ Pichia Expression Kit Manual (Invitrogen). Transformants were selected on YPD plates containing different concentrations of Zeocin (100-2000 μg/mL) and incubated at 30°C for 72 hours. Individual colonies were screened for expression under induction with methanol (0.5-2%) at the 10 mL culture level (BMMY medium) as described^12, 22^. The expression level of select colonies was identified by SDS-PAGE and immunoblotting using anti-SARS-CoV-2 antibodies (anti-SARS-CoV-2 spike rabbit monoclonal antibody, Sino Biological, Cat # 40150-R007), and research seed stocks of the highest expressing clones were frozen at −80 °C.

RBD219-WT and RBD219-N1C1 were expressed at the 5 L scale using a Celligen 310 benchtop fermentation system (Eppendorf). For the RBD-WT, 2.5 L of basal salt medium were inoculated with a seed culture to an initial OD_600_ of 0.05 and grown at 30 °C, pH 5.0 with 30% dissolved oxygen until glycerol depletion. During the first hour of methanol induction, the temperature was reduced from 30 °C to 26 °C and the pH was increased from 5.0 to 6.0. After approximately 70 hours of induction (methanol feed at 1-11 mL/L/hr), the culture was harvested from the fermenter, and cells were removed by centrifugation for 30 min at 12,227 x g at 4 °C. For RBD219-N1C1, the fermentation process was slightly different in that low salt medium was used, the induction temperature was set to 25 °C and the pH to 6.5 and, the methanol feed rate was between 1-15 ml/L/hr. The fermentation supernatant (FS) was filtered (0.45 μm PES filter) and stored at −80 °C before purification.

A hexahistidine-tagged SARS-CoV-2 RBD219-WT was purified from fermentation supernatant (FS) by immobilized metal affinity chromatography followed by size exclusion chromatography (SEC). The FS was concentrated, and buffer exchanged to buffer A (20 mM Tris-HCl pH 7.5 and 0.5 M NaCl) using a Pellicon 2 cassette with a 10 kDa MWCO membrane (MilliporeSigma) before being applied to a Ni-Sepharose column (Cytiva). The column was washed with buffer A plus 30 mM imidazole and elution was undertaken in buffer A containing 250 mM imidazole. The RBD219-WT protein was further purified using a Superdex 75 prep grade column (Cytiva) pre-equilibrated in buffer B (20 mM Tris-HCl pH 7.5 and 150 mM NaCl) after concentrating eluates from the Ni column using an Amicon Ultra-15 concentrator with a 10 kDa MWCO membrane (MilliporeSigma). Monomeric RBD219-WT was pooled, aseptically filtered using a 0.22 μm filter, and stored at −80 °C.

For the purification of the tag-less RBD219-N1C1, ammonium sulfate salt was added to the FS to a final concentration of 1 M (w/v) before the sample was applied to a Butyl Sepharose HP column (Cytiva). The column was washed with buffer C (30 mM Tris-HCl pH 8.0) with 1 M ammonium sulfate and protein was eluted with buffer C containing 0.4 M ammonium sulfate. A polishing step using a Superdex 75 prep grade column (Cytiva) pre-equilibrated in buffer B followed.

### 2.2 SDS-PAGE

To evaluate the size of RBD219-WT and RBD219-N1C1, 2 μg of these two proteins were loaded onto a 4-20% tris-glycine gel under non-reduced and reduced conditions. These two proteins were also treated with PNGase-F (NEB) under the reduced condition to remove N-glycans and loaded on the gel to assess the impact of the glycans on the protein size. Gels were stained using Coomassie Blue and analyzed using a Bio-Rad G900 densitometer with Image Lab software.

### 2.3 Vaccine formulation and Alhydrogel^®^-protein binding study

SARS-CoV-2 RBD219-N1C1 was diluted in 20 mM Tris, 150 mM NaCl, pH 7.5 (TBS buffer) before mixing with Alhydrogel^®^ (aluminum oxy-hydroxide; Catalog # 250-843261 EP, Brenntag). To calculate the Langmuir binding isotherm of RBD219-N1C1 to Alhydrogel^®^, RBD219-N1C1 and Alhydrogel^®^ were mixed at different ratios (from 1:2 to 1:20). The RBD219-N1C1/Alhydrogel^®^ mixture was stored for one hour at RT, to reach an equilibrium state. The Alhydrogel^®^ formulations were centrifuged at 13,000 x g for 5 min, and the supernatant was removed. The protein in the supernatant fraction and the pellet fraction were quantified using a micro-BCA assay (ThermoFisher).

### 2.4 ACE-2 binding assay

For the ACE-2 binding study, the Alhydrogel^®^-RBD vaccine formulations were blocked overnight with 0.1% BSA. After hACE-2-Fc (LakePharma) was added, the samples were incubated for 2 hours at RT. After incubation, the Alhydrogel^®^ was spun down at 13,000 x g for 5 min. The hACE-2-Fc which did not bind to the RBD on the Alhydrogel^®^ remained in the supernatant. The hACE-2-Fc content in the supernatant was quantified by ELISA using 96-Well MaxiSorp Immuno plates (ThermoFisher) coated overnight with 200 ng/well of RBD219-WT protein. After blocking with 0.1% BSA, 100 μL supernatant samples were added to each well. Plates were washed 4 times with an automated plate washer using PBS with Tween (PBST). A secondary antibody against human Fc was used to detect hACE-2-Fc bound the proteins on the plate. Plates were washed 5 times with an automated plate washer using PBST before 100 μL TMB solution were added. The enzymatic reaction was stopped with HCl and absorption readings were made at 450 nm. The final concentration of the hACE-2 bound on the Alhydrogel^®^ was determined as [hACE-2-Fc on Alhydrogel^®^] = [Total hACE-2-Fc] – [hACE-2-Fc in supernatant].

### 2.5 Immunogenicity testing

To examine RBD-specific antibodies in mouse sera, indirect ELISAs were conducted. 96-well NUNC ELISA plates were coated with 2 μg/mL RBD219-WT in 100 μL 1x coating buffer per well and incubated overnight at 4 °C. The next day the coating buffer was discarded, and plates were blocked with 200 μL/well 0.1% BSA in PBST for 2 hours at room temperature. Mouse serum samples were diluted from 1:200 to 1: 437,400 in 0.1% BSA in PBST. Blocked ELISA plates were washed once with 300 μL PBST using a Biotek 405TS plate washer and diluted mouse serum samples were added to the plate in duplicate, 100 μL/well. As negative controls, pooled naïve mouse serum (1:200 diluted) and blanks (0.1% BSA PBST) were added as well. Plates were incubated for 2 hours at room temperature, before being were washed four times with PBST. Subsequently, 1:6,000 diluted goat anti-mouse IgG HRP antibody (100 μL/well) was added in 0.1% BSA in PBST. Plates were incubated for 1 hour at room temperature, before washing five times with PBST, followed by the addition of 100 μL/well TMB substrate. Plates were incubated for 15 min at room temperature while protected from light. After incubation, the reaction was stopped by adding 100 μL/well 1 M HCl. The absorbance at a wavelength of 450 nm was measured using a BioTek Epoch 2 spectrophotometer. Duplicate values of raw data from the OD_450_ were averaged. The titer cutoff value was calculated using the following formula: Titer cutoff = 3 x average of negative control + 3 x standard deviation of the negative control. For each sample, the titer was determined as the lowest dilution of each mouse sample with an average OD_450_ value above the titer cutoff. When a serum sample did not show any signal at all and a titer could not be calculated, an arbitrary baseline titer value of 67 was assigned to that sample (baseline).

### 2.6 Pseudovirus assay

Pseudovirus was prepared in HEK-293T cells by previously reported methods with modifications^24^. Cells were transfected with 2.5 μg of the plasmid encoding the SARS-CoV-2 spike protein (p278-1^25^) and 3.7 μg of luciferase-encoding reporter plasmid (pNL4-3.lucR-E^26^) and Gag/Pol-encoding packaging construct (pΔ8.9^27^). Pseudovirus containing supernatant was recovered after 48◻h and passed through a 0.45 μM filter before use.

For each serum sample, 30 μL pseudovirus were incubated with serial dilutions of heat-inactivated serum (eight dilutions in a 4-fold stepwise manner) for 1 h at 37 °C. Next, 100 μL of these sera-pseudovirus mixtures were added to 293T-hACE2 cells in 96-well poly-D-lysine coated culture plates. Following 48 h incubation in a 5% CO2 environment at 37 °C, the cells were lysed with 100 μL Promega Glo Lysis buffer, 15 min RT. Finally, 20 μL lysate was added to 100 μL Luc substrate (Promega Luciferase Assay System). The amount of luciferase was quantified by luminescence (relative luminescence units (RLU)), using a Promega GloMax luminometer (Steady-Glo program). The percent virus inhibition was calculated as (1-RLU of sample/ RLU of negative control) x 100. Serum from vaccinated mice was also characterized by the IC50-value, defined as the serum dilution at which the virus infection was reduced to 50% compared with the negative control (virus◻+◻cells). When a serum sample did not neutralize 50% of the virus when added at a 1:10 dilution, the IC50 titer could not be calculated and an arbitrary baseline titer value of 10 was assigned to that sample (baseline). As a control, human convalescent sera for SARS-CoV-2 (NIBSC 20/130) were used (National Institute for Biological Standards and Control).

### 2.7 Cytokine analysis

#### 2.7.1 Preparation of splenocytes for restimulation

Single-cell suspensions from mouse splenocytes were prepared using a cell dissociator (GentleMACS Octo Dissociator, Miltenyi Biotec) based on a previously optimized protocol^28^. The concentration and the viability of the splenocyte suspensions were measured after mixing with AOPI dye and counted using the Nexcelom Cellometer Auto 2000.

For the re-stimulation assays, splenocyte suspensions were diluted to 8×10^6^ live cells/mL in a 2-mL deep-well dilution plate and 125 μL of each sample was seeded in two 96-well tissue culture treated culture plates. Splenocytes were re-stimulated with 10 μg/mL RBD219-WT, 20 ng/mL PMA + 1 μg/mL Ionomycin or just media (unstimulated). For the flow cytometry plate, the PMA/I was not added until the next day. 125 μL (2x concentration) of each stimulant was mixed with the 125 μL splenocytes suspension in the designated wells. After all the wells were prepared, the plates were incubated at 37 °C 5% CO_2_. One plate was used for the cytokine release assay, while the other plate was used for flow cytometry. For flow cytometry, another plate was prepared with splenocytes, which would be later used as fluorescence minus one – controls (FMOs).

#### 2.7.2 In vitro *cytokine release assay*

After 48 hours in the incubator, splenocytes were briefly mixed by pipetting. Then plates were centrifuged for 5 min at 400 x g at RT. Without disturbing the pellet 50 μL supernatant was transferred to two skirted PCR plates and frozen at – 20 °C until use.

For the *in vitro* cytokine release assay, splenocytes were seeded in a 96-well culture plate at 1×10^6^ live cells in 250 μL cRPMI. Splenocytes were then (re-)stimulated with either 10 μg/mL RBD219-WT protein, 10 μg/mL RBD219-N1C1 protein, PMA/I (positive control), or nothing (negative control) for 48 hours at 37 °C 5% CO_2_. After incubation, 96-well plates were centrifuged to pellet the splenocytes down and supernatant was transferred to a new 96-well plate. The supernatant was stored at −20°C until assayed. A Milliplex Mouse Th17 Luminex kit (MD MilliPore) with analytes IL-1β, IL-2, IL-4, IL-6, IL-10, IL-12(p70), IL-13, IL-17A, IL-23, IFN-γ, and TNF-α was used to quantify the cytokines secreted in the supernatant by the re-stimulated splenocytes. An adjusted protocol based on the manufacturers’ recommendations was used with adjustments to use less sample and kit materials^29^. The readout was performed using a MagPix Luminex instrument. Raw data was analyzed using Bio-Plex Manager software, and further analysis was done with Excel and Prism.

#### 2.7.3 Cytokine production of activated CD4+ and CD8+ T cells

Surface staining and intracellular cytokine staining followed by flow cytometry were performed to measure the amount of activated (CD44+) CD4+ and CD8+ T cells producing IFN-γ, IL-2, TNF-α, and IL-4 upon re-stimulation with S2RBD219 WT.

Five hours before the 24-hour re-stimulation incubation, Brefeldin A was added to block cytokines from secretion. PMA/I was also added to designated wells as a positive control. After the incubation, splenocytes were stained for the relevant markers. A viability dye and an Fc Block were also used to remove dead cells in the analysis and to minimize non-specific staining, respectively.

After staining, splenocytes were analyzed using an Attune NxT flow cytometer instrument at the Baylor College of Medicine Cytometry and Cell Sorting Core. Raw data was analyzed in VenturiOne software and gating results were copied in Excel. The %-gating values from the non-stimulated controls were subtracted from the re-stimulated controls to observe the difference in %-gating induced by the re-stimulation.

The gating strategy from the analysis of the results is shown in **Supplemental Figure 1.** From all events collected the doublets are removed to obtain only single-cell events. Then events are selected on size and granularity to obtain splenocytes only. Following the removal of dead splenocytes, a gate is set to only select activated (CD44+) T cells (CD3+)^30^. T cells are then separated into CD4+ T helper cells and CD8+ cytotoxic T cells. For T helper cells the events positive for IFN-γ, TNF-α, IL-2, and IL-4 were selected, while for cytotoxic T cells only IFN-γ, TNF-α, and IL-2 positive events were gated.

### 2.8 Statistical analysis

To test for significant differences between groups in ELISA, Luminex, and flow cytometry results, Kruskal-Wallis tests in combination with Dunn’s multiple comparison tests were performed. ns (not significant): p>0.05, *: p < 0.05 and **: p < 0.01.

## 3. Results

Here we report on the expression of a modified, recombinant RBD of the SARS-CoV-2 spike protein using a yeast (*P. pastoris)* expression system. The candidate antigen selection, modifications, and production processes were based on eight years of process development, manufacture, and preclinical prior experience with a SARS-CoV recombinant protein-based receptor-binding domain (RBD)^11–13^. The RBDs of SARS-CoV-2 and SARS◻CoV share significant amino acid sequence similarity (>75% identity, >80% homology) and both use the human angiotensin-converting enzyme 2 (ACE2) receptor for cell entry^31, 32^. Process development using the same procedures and strategies used for the production, scale-up, and manufacture of the SARS-CoV recombinant protein allowed for a rapid acceleration in the development of a scalable and reproducible production process for the SARS◻CoV-2 RBD219-N1C1 protein, suitable for technological transfer to a manufacturer.

We found that the modifications used to minimize yeast-derived hyperglycosylation and optimize yield, purity, and stability of the SARS-CoV RBD219-N1 protein were also relevant to the SARS-CoV-2 RBD expression and production process. The modified SARS-CoV-2 antigen, RBD219-N1C1, when formulated on Alhydrogel^®^, was shown to induce virus-neutralizing antibodies in mice, equivalent to those levels elicited by the wild-type (RBD219-WT) recombinant protein counterpart.

### 3.1 Cloning and expression of the modified SARS-CoV-2 RBD

The wild-type SARS-CoV-2 RBD amino acid sequence comprises residues 331-549 of the spike (S) protein (GenBank: QHD43416.1) of the Wuhan-Hu-1 isolate (GenBank: MN908947.3) ( **Legends:** Figure 1). In the RBD-219-WT construct, the gene fragment was expressed in *P. pastoris*. After fermentation at the 5 L scale, the hexahistidine-tagged protein was purified by immobilized metal affinity chromatography, followed by size-exclusion chromatography. We observed glycosylation and aggregation during these initial expression and purification studies, and therefore, similar to our previous strategy^12^, we generated a modified construct, RBD219-N1C1, by deleting the N331 residue and mutating the C538 residue to alanine. The additional mutation of C538 to A538 was done because we observed that in the wild-type sequence nine cysteine residues likely would form four disulfide bonds. Therefore, the C538 residue was likely available for intermolecular cross-linking, leading to aggregation. As a result, in the RBD219-N1C1 construct, *Pichia*-derived hyperglycosylation, as well as aggregation via intermolecular disulfide bridging, were greatly reduced^21^. We note that the deleted and mutated residues are structurally far from the immunogenic epitopes and specifically the receptor-binding motif (RBM) of the RBD ( **Legends:** Figure 1). On SDS-PAGE tris-glycine gels, the RBD219-WT protein migrated at approximately 28 kDa under non-reduced conditions and 33 kDa under reduced condition, while the RBD219-N1C1 protein migrated at approximately 24 kDa under non-reduced condition and 29 kDa under reduced condition. However, after N-glycans were removed enzymatically, these two proteins showed a similar molecular weight of approximately 25 kDa (**Supplemental Figure 2**). The purity of both proteins was analyzed by densitometry showing levels of >95%.

**Figure 1.**
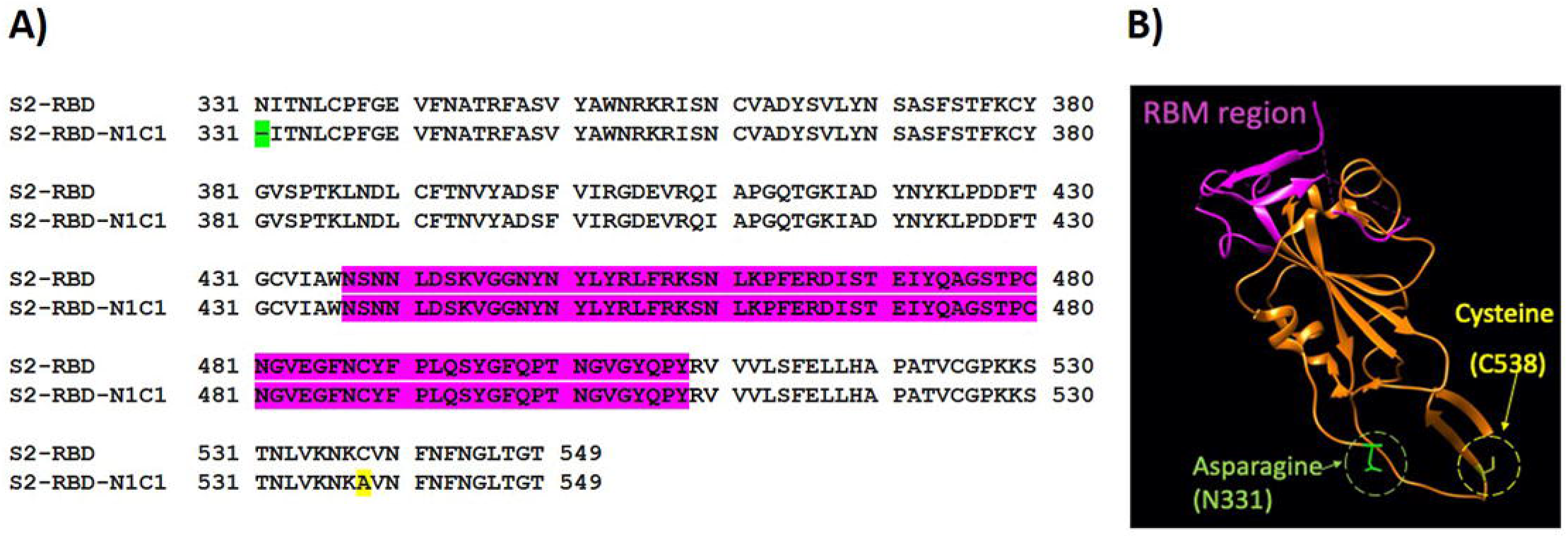
A) Amino acid sequence alignment between SARS-CoV-2 RBD219-WT (S2-RBD) and RBD219-N1C1 (S2-RBD-N1C1). In the N1C1-mutant, the N-terminal glutamine residue (N331, green) is removed and a C538A mutation (yellow) was introduced. Neither mutation is inside the receptor-binding motif (RBM, purple). B) The structure model of RBD219-WT was extracted from the crystal structure of the SARS-CoV-2 spike protein (PDB ID 6VXX). The RBM (N436-Y508) is again shown in purple while the deleted asparagine (N331) and mutated cysteine (C538, mutated to alanine) in RBD219-N1C1 are highlighted in green and yellow, respectively.

### 3.2 ACE-2 binds to recombinant SARS◻CoV-2 RBD219-N1C1 protein formulated on Alhydrogel^®^

When mixing 25 μg of either RBD219-WT or RBD219-N1C1 proteins with 500 μg of Alhydrogel^®^, we observed that >98% of the proteins bound to Alhydrogel^®^ after 15 min of incubation. Only when the Alhydrogel^®^ was reduced to less than 100 μg (Alhydrogel^®^/RBD219 ratio <4), the Alhydrogel^®^ surface was saturated, and protein started to be detected in the supernatant (**Figure 2A)**. It is known that unbound protein may impact the immunogenicity of the vaccine formulation, therefore we proceeded to only evaluate formulations with Alhydrogel^®^/RBD219 ratios higher than 4.

**Figure 2.**
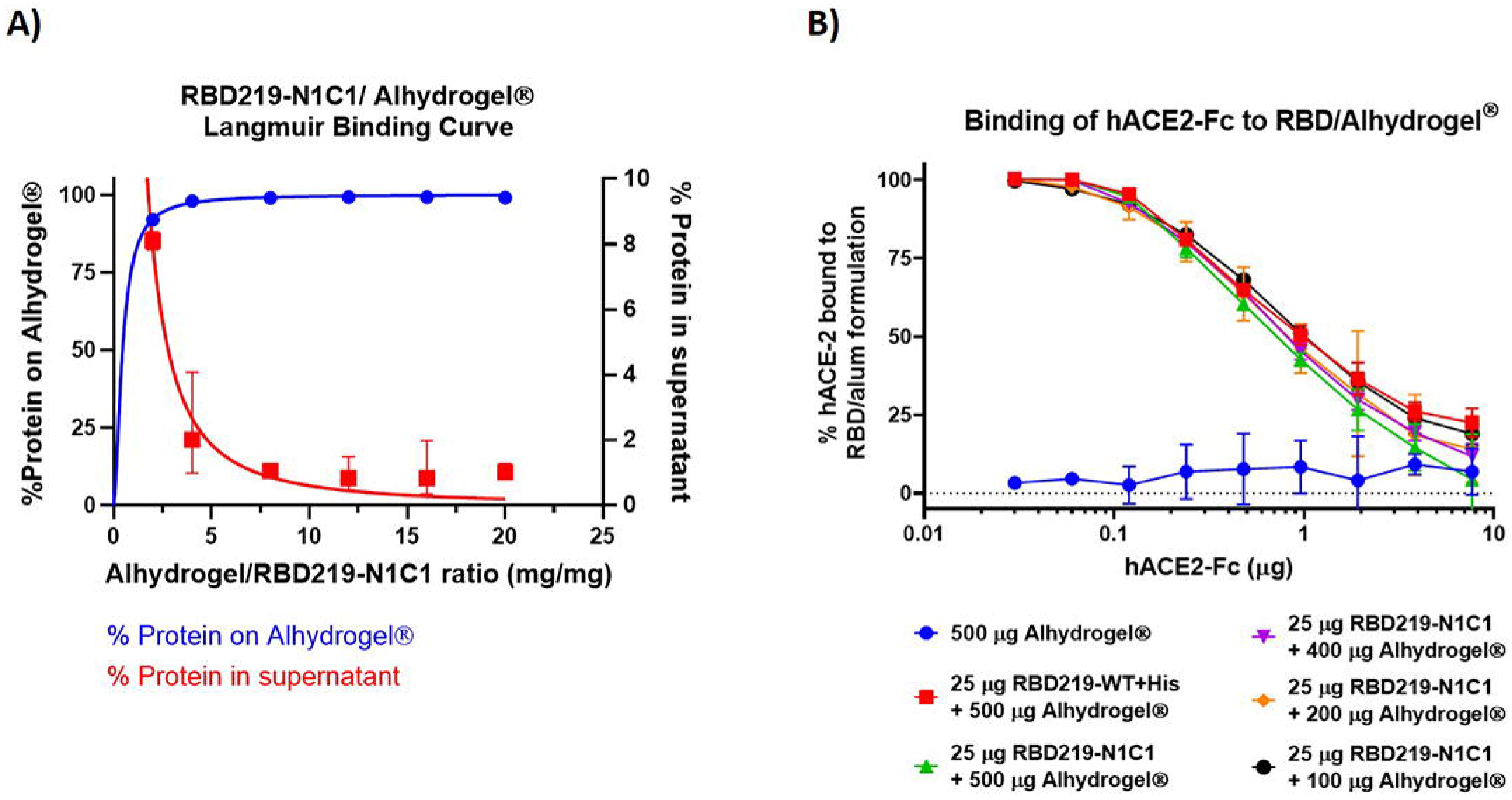
A) Langmuir binding isotherm of RBD219-N1C1 to Alhydrogel^®^. B) ELISA data, comparing the binding interaction of hACE-2-Fc to RBD219-WT bound Alhydrogel^®^ (red) and RBD219-N1C1 bound on different amounts of Alhydrogel^®^ (green, purple, orange, and black). Five hundred μg Alhydrogel^®^ alone served as a negative control (blue). Data are shown as the geometric mean (n=3) with 95% confidence intervals.

**Figure 2B** shows that hACE-2-Fc, a recombinant version of the human receptor used by the virus to enter the host cells, can bind with the RBD proteins that are adsorbed on the surface of the Alhydrogel^®^ This demonstrates that bound RBD proteins are structurally and possibly functionally active and that after adsorption the protein does not undergo any significant conformational changes that could result in the loss of possible key epitopes around the receptor-binding motif (RBM). The finding was consistent with the ACE-2 binding assay performed on the RBD proteins in the ELISA plate without being pre-adsorbed to Alhydrogel^®21^.

We saw no statistical differences between the binding of hACE-2-Fc to RBD219-WT (red, Figure 2B) or RBD219-N1C1 (green, Figure 2B) proteins, based on an unpaired t-test (P=0.670). Likewise, we saw no relation between the amount of Alhydrogel^®^ to which the RBD was bound and the interaction with hACE-2-Fc, indicating that the surface density of the RBD proteins on the Alhydrogel^®^ plays no role in the presentation of ACE binding sites.

### 3.3 Recombinant RBD219-N1C1 protein, formulated with Alhydrogel^®^, elicits a strong neutralizing antibody response in mice

Recombinant RBD219-N1C1 protein (25 μg) was formulated with various amounts (100 – 500 μg) of Alhydrogel^®^. Controls included a cohort receiving only Alhydrogel^®^ and another receiving the RBD219-WT antigen, also formulated with 500 μg Alhydrogel^®^. Six- to eight-week-old female BALB/c mice were immunized 2-3 times subcutaneously at approximately 21-day intervals (**Figure 3A**). Blood samples were taken on day 35 from all study animals to assess total IgG antibody titers, as well as neutralizing antibody titers (Dataset 1). Half of the mice, those with the highest IgG titers in their respective group, were euthanized on day 43 to allow the evaluation of the cellular immune response after two immunizations. For this dataset (Dataset 2), we also measured IgG and neutralizing antibody titers. The remaining mice received a third vaccination on day 43 and were euthanized on day 57 for the assessment of humoral and cellular immune responses (Dataset 3).

**Figure 3.**
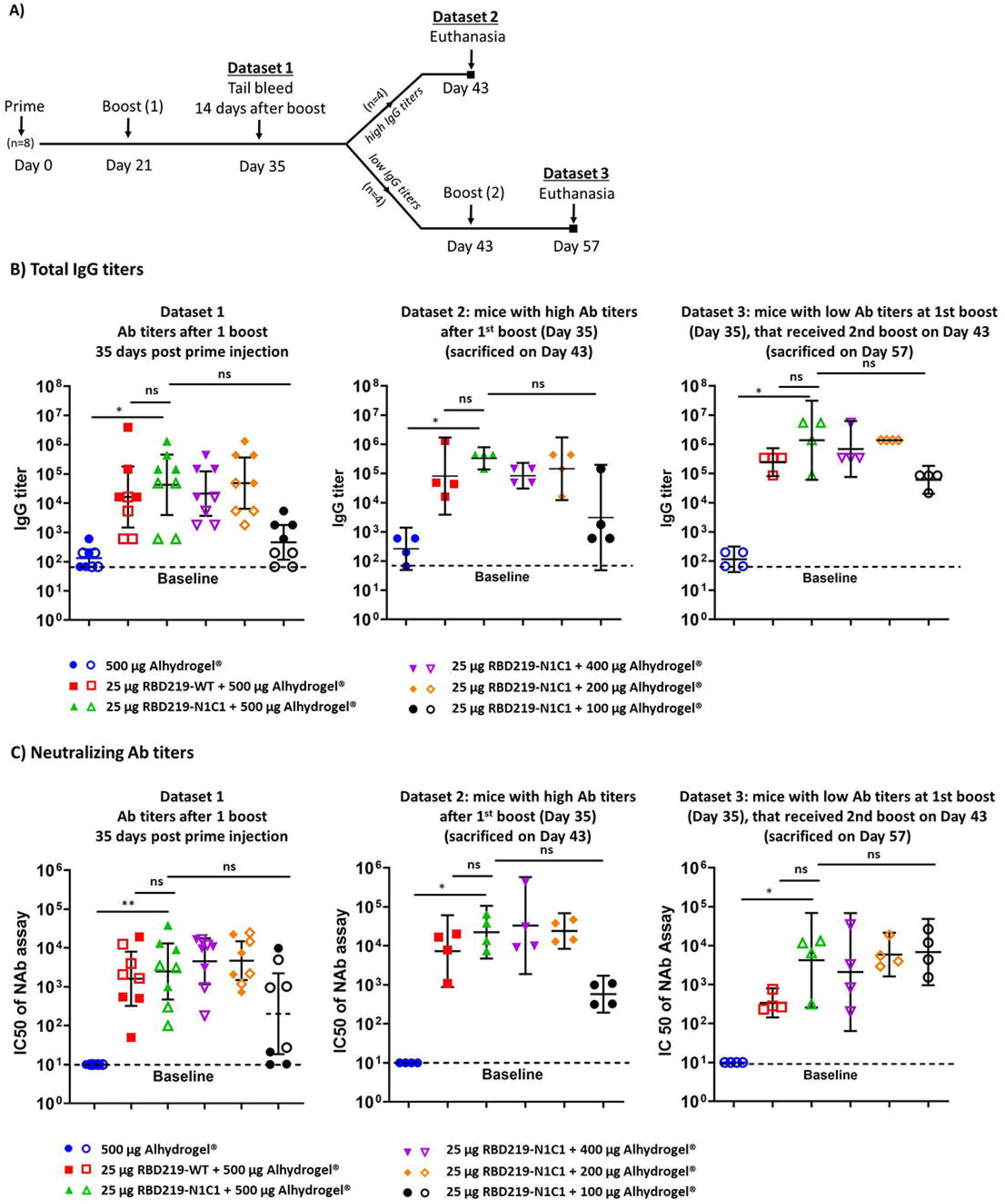
A) Study design. B) Total IgG titers of Datasets 1, 2, and 3 measured respectively at days 35, 43, and 57 post the prime injection. IgG titers were determined against RBD219-WT protein. Closed data points represent data from mice with the highest IgG titers (Dataset 2), open data points represent data from mice with the lowest IgG titers (Dataset 3). C) IC50 values measured by a pseudovirus neutralization assay. Datasets 1, 2, and 3 are measured respectively at day 35, 43, and 57 after the first injection. Baselines indicate the lowest dilution measured. Lines on each group represent the geometric mean and 95% confidence intervals.

#### Humoral immune response

On day 35 (Dataset 1), after receiving two vaccinations, all groups that had received the recombinant protein formulated with at least 200 μg Alhydrogel^®^ produced similar and robust IgG titers. The group receiving the protein with only 100 μg Alhydrogel^®^, produced a lower IgG response, albeit slightly higher than the negative control that had been immunized with 500 μg Alhydrogel^®^ alone (**Figure 3B, Supplemental Table 1**). Importantly, based on a Mann-Whitney test, we determined that there was no statistical difference between the groups vaccinated with the modified and the wild-type version of the RBD protein (p=0.3497). The average neutralizing antibody titers observed on day 35 (IC50 range: 5.0×10^3^ to 9.4×10^3^, **Supplemental Table 2**) matched with the total IgG titers, showing equally high IC50 values for all vaccines that contained at least 200 μg Alhydrogel^®^ and lower IC50 values for the vaccine with only 100 μg Alhydrogel^®^ and no IC50 values for the adjuvant-only control (**Figure 3C**).

On day 43, 22 days after receiving the boost vaccination, half of the mice in each group (N=4), those with the highest IgG titers, were euthanized to determine the total IgG, the IgG subtypes, and the neutralizing antibody titers. As we observed on day 35, all animals that had received the vaccine produced strong antibody titers, with the groups receiving ≥200 μg Alhydrogel^®^ eliciting a higher titer than those that received only 100 μg of Alhydrogel^®^, albeit no statistical significance was detected (**Figures 3B**). For all animals, as typical for vaccine formulations containing aluminum, the IgG2a:IgG1 titer ratio was <0.1 (**Supplemental Figure 3).** In the pseudovirus neutralization assay for the day 43 samples (**Figure 3C**), all vaccines containing ≥200 μg Alhydrogel^®^ elicited IC50 titers that, on average, were several-fold higher than on day 35 (IC50 range: 1.1×10^4^ to 1.2×10^5^, **Supplemental Table 2**). There again was no difference between the RBD219-WT and RBD-N1C1 vaccines.

On day 57, all remaining animals were euthanized. In contrast to the animals studied on days 35 and 43, these animals had received a second boost vaccination. A robust immune response in all vaccinated mice, including those immunized with the protein adsorbed to 100 μg Alhydrogel^®^ achieved high average IgG titers. The total IgG titers in the mice euthanized on day 57, had increased after the third vaccination, compared to the titers seen on day 35. Likewise, we observed a corresponding increase in the average IC50 values (IC50 range: 3.8×10^2^ to 1.1×10^4^, **Supplemental Table 2**) for all animals, including those immunized with the protein adsorbed to 100 μg Alhydrogel^®^. Interestingly, for this time point, the cohort receiving 25 μg RBD219-N1C1 with 500 μg Alhydrogel^®^ appeared to show higher neutralizing antibody titers than the corresponding RBD219-WT group, albeit that difference was not statistically significant.

#### Cellular immune response

For all animals euthanized on day 43 (having received two vaccinations) and day 57 (having received three vaccinations), the cellular immune response was characterized through the restimulation of isolated mouse splenocytes with the recombinant RBD219-WT protein. For all samples, we employed Flow Cytometry to quantify intracellular cytokines in CD4+ and CD8^+^ cells after restimulation (**Figure 4A**). On day 43, high percentages of CD4^+^-IL-4 and, to a slightly lesser extent CD4^+^-TNFα producing cells were detected. Conversely, as expected for an Alhydrogel^®^-adjuvanted vaccine, low levels of IL-2 producing CD4^+^ cells were seen. In a cytokine release assay, strong IFN-γ, IL-6, and IL-10 secretion was observed independent of whether the animals had received two or three immunizations, whereas low amounts of secreted Th1-typical cytokines such as IL-2 or IL-12 were seen (**Figure 4B**).

**Figure 4:**
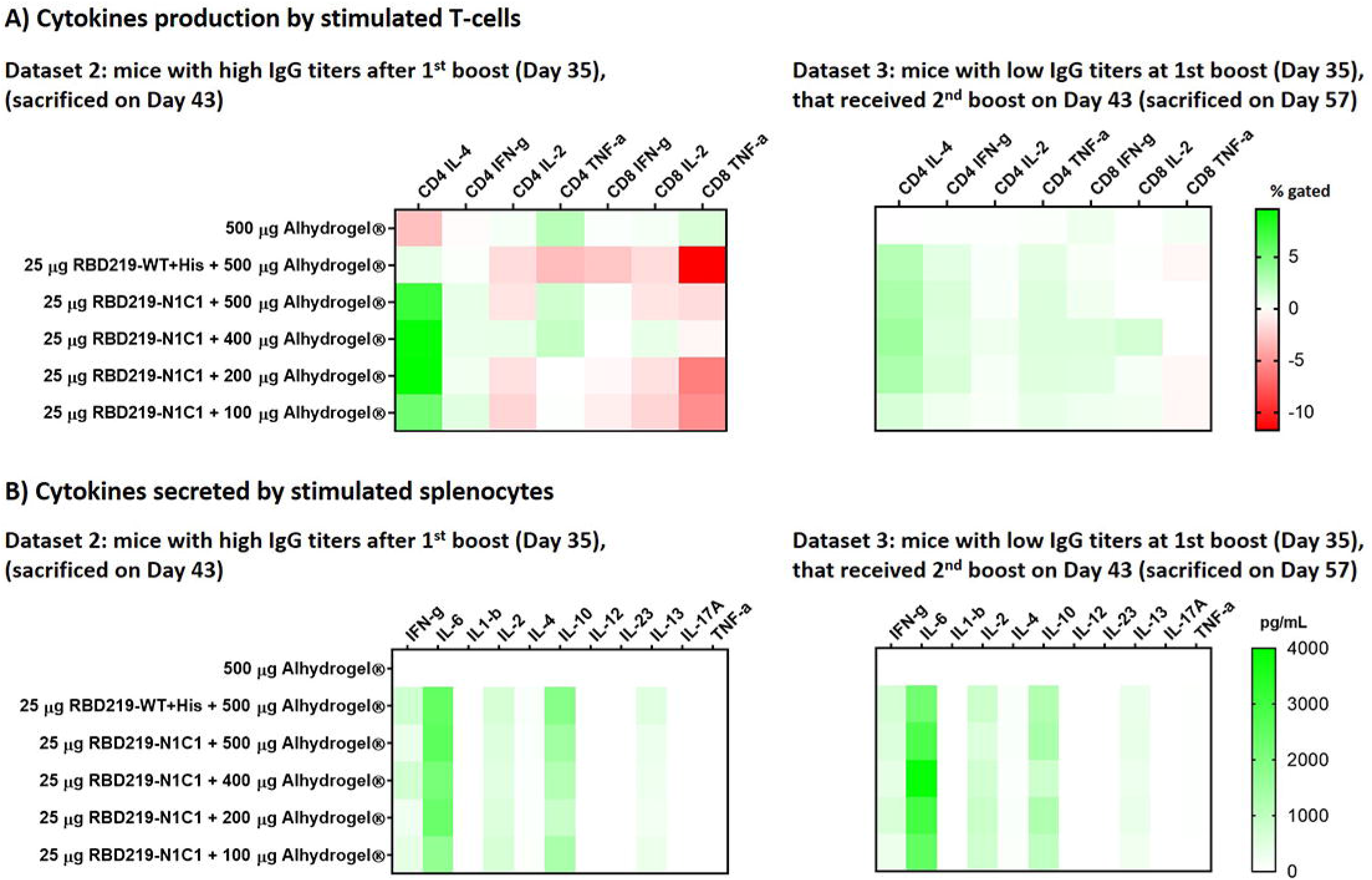
A) Heatmap of the cytokine response of CD4+ and CD8+ T cells after restimulation with SARS-CoV-2 RBD219-WT or RBD219-N1C1, re-stimulated splenocytes were surface and intracellularly stained and subsequently analyzed by flow cytometry. Splenocytes were obtained from mice who received two vaccinations (day 43) or three vaccinations (Day 57). Non-stimulated controls were subtracted from re-stimulated samples. B) Heatmap of secreted cytokines in supernatant from re-stimulated splenocytes from mice who received two vaccinations (day 43) or three vaccinations (Day 57). Cytokine concentrations of non-stimulated controls were subtracted from re-stimulated samples.

### 3.4 Study limitations

The authors recognize that the data presented here has limitations. For instance, we do not include viral challenge studies. Prediction of the vaccine’s *in vivo* efficacy is therefore based on the neutralizing activity of the vaccine-induced antibodies in mice, measured through an *in vitro* pseudovirus assay. A challenge study in non-human primates is currently ongoing and will be published separately. Moreover, our studies were limited to aluminum-based formulations. Further studies are needed to investigate the combination of the vaccine antigen with different adjuvant and immunostimulant formulations, which may lead to dose spearing, reduce the number of doses, and/or induce a more balanced and long-lasting immune response.

## 4. Discussion

Here we report on a yeast-expressed SARS-CoV-2 RBD219-N1C1 protein and its potential as a vaccine antigen candidate for preventing COVID-19. Building on extensive prior experience developing vaccines against SARS-CoV and MERS-CoV^12–14^, we selected and compared the SARS-CoV-2 RBD219-WT and the SARS-CoV-2 RBD219-N1C1 proteins for their potential to induce high titers of virus-neutralizing antibodies, T-cell responses, and protective immunity. The decision to focus our antigen development on the RBD of the viral spike protein stems in part from earlier work on the ancestral SARS vaccine, where we had already been able to demonstrate low-cost, high-yield production of that homologous vaccine antigen^33^, something that we now also confirmed formally for the SARS-CoV-2 RBD antigen^34^. In addition, the original SARS-CoV RBD-based vaccine was superior to the full-length spike protein at inducing specific antibodies and fully protected mice from SARS-CoV infection while preventing eosinophilic pulmonary infiltrates in the lungs upon challenge^11^, a process more likely linked to Th17-dominant and mixed Th1/Th2 responses, when compared to using full-length spike protein as the antigen^35^. As an additional argument in favor of the RBD, Liu *et al.* have reported that most neutralizing epitopes of the SARS-CoV-2 spike protein are in the RBD and the N-terminal domain (NTD), with the most potent ones near the RBD’s Receptor Binding Motif^36^.

While it is known that yeast N-glycosylation is different from that in mammalian cells and yeast-derived glycosylation of the eight N-glycosylation sites in the NTD might alter the induction of neutralizing antibodies, little impact is expected for the RBD since there are no glycosylation sites near the RBM. Moreover, in this work, using the SARS-CoV-2 RBD219 protein homolog, we observed that, just like in the case of the SARS-CoV RBD antigen, the deletion of the N-terminal asparagine residue reduced hyperglycosylation, thus allowing for easier purification of the antigen obtained from the yeast expression system. This finding was confirmed when comparing the minimal changes noted in apparent molecular weights of our reduced RBD compared to an enzymatically deglycosylated form. As an additional modification, mutagenesis of a free cysteine residue further improved protein production through the reduction of aggregation. Based on the predicted structure of the RBD, no impact on the functionality of the RBD219-N1C1 antigen was expected, and using an ACE-2 *in vitro* binding assay we indeed showed similarity to the RBD219-WT antigen. In addition, we showed that, in mice, the modified RBD219-N1C1 antigen triggered an equivalent immune response to the RBD219-WT protein when both proteins were adjuvanted with Alhydrogel^®^.

Similar to our previous findings with the SARS-CoV RBD antigen^11^, we show that RBD219-N1C1 when formulated with Alhydrogel^®^ elicits a robust neutralizing antibody response with IC50 values up to 4.3×10^5^ in mice, as well as an expected T-cell immunological profile. Some of the titers of virus-neutralizing antibodies exceed the titer, 2.4×10^4^, measured in-house with human convalescent serum research reagent for SARS-CoV-2 (NIBSC 20/130, National Institute for Biological Standards and Control, UK). Among the T-cell activation markers, we found high percentages of CD4+ T cells expressing IL-4 and IFN-γ, possibly indicating increased numbers of functional T follicular helper cells which will support the generation of antibody-producing plasma cells and long-lived memory B cells^37^.

In a mouse virus challenge model for the SARS-CoV RBD recombinant protein vaccine, we found that Alhydrogel^®^ formulations induced high levels of protective immunity but did not stimulate eosinophilic immune enhancement, suggesting that Alhydrogel^®^ may even reduce immune enhancement. This prior experience offers the potential for Alhydrogel^®^ as a key adjuvant for consideration during coronavirus vaccine development^38^. Such findings have led to a reframing of the basis for immune enhancement linked to coronavirus respiratory infections^35^. A recent analysis and review by the NIH ACTIV Vaccine Working Group confirmed that aluminum or Th2 responses remain viable options for vaccine development concluding that “it is not possible to clearly prioritize or down-select vaccine antigens, adjuvants, biotechnology platforms, or delivery mechanisms based on general immunological principles or the available preclinical data”^39^.

Therefore, the RBD219-N1C1 vaccine antigen on Alhydrogel^®^ merits its evaluation as a COVID-19 vaccine with or without other immunostimulants. Looking at the landscape of recombinant protein-based COVID-19 vaccines, the WHO lists several advanced COVID-19 candidates that are based on recombinant proteins^1^, and at least seven COVID-19 vaccines include aluminum as part of the adjuvant component^8, 15, 17, 18, 40–46^, often in combination with other immunostimulants, such as CpG, to achieve a balanced immune response. These recombinant protein vaccines, including RBD219-N1C1, might find an additional important use as a booster if one of the newer platform vaccines, e.g., mRNA or adenovirus-based vaccines induce lower than expected immunogenicity or protection. Likewise, we see an opportunity for protein vaccines representing the newly appearing SARS-CoV-2 variants^47^ to act as boosters in individuals previously immunized with the wild-type antigen. Such prime-boost approaches have been used successfully with the chimp adenovirus vaccine for malaria and other systems^48, 49^, and are currently being entertained for SARS-CoV-2^50^.

The selection of the *P. pastoris* expression platform for the production of the RBD antigen was motivated by the intent to develop a low-cost production process that could easily be transferred to manufacturers in LMICs. Currently, there are several types of COVID-19 vaccine candidates in advanced clinical trials^7, 51–56^ that, as they require advanced infrastructure, focus on the developed world. Being able to match the existing experience in LMICs with the production of other biologics in yeast increases the probability of successful technology transfer. For example, currently, the recombinant hepatitis B vaccine is produced in yeast by several members of the Development Country Vaccine Manufacturers Network (DCVMN), and we foresee that, given the existing infrastructure and expertise, those facilities could be repurposed to produce a yeast-produced COVID-19 vaccine^58^. Recently, the research cell bank and the production process for the RBD219-N1C1 antigen were technologically transferred to a vaccine manufacturer in India and produced under cGMP conditions; adjuvanted with Alum and CpG and that vaccine is currently in clinical trials in India^59^. In parallel, studies to adapt the RBD219-WT and RBD219-N1C1 antigens to the recently appearing SARS-CoV-2 variants are underway, as is a SARS-CoV-2 challenge study in a non-human primate model. Given the limited access to suitable test facilities, the onset of that study was significantly delayed, but we expect to publish its results soon.

## Acknowledgments

This work was supported by the Robert J. Kleberg Jr. and Helen C. Kleberg Foundation, as well as the NIH (AI14087201), Baylor College of Medicine, and Texas Children’s Hospital. This project was further supported by the Cytometry and Cell Sorting Core at Baylor College of Medicine with funding from the CPRIT Core Facility Support Award (CPRIT-RP180672), the NIH (CA125123 and RR024574), and the assistance of Joel M. Sederstrom. We are grateful to Dr. Vincent Munster (NIAID) for providing the spike expression plasmid for SARS-CoV-2.

