## Supplemental for "SARS-CoV-2 RBD219-N1C1: A Yeast-Expressed SARS-CoV-2 Recombinant Receptor-Binding Domain Candidate Vaccine Stimulates Virus Neutralizing Antibodies and T-cell Immunity in Mice"

### 1 Supplemental Methods

### 1.1. SDS-PAGE

### To evaluate the size of RBD219-WT and RBD219-N1C1, 2 μg of these two proteins were loaded onto a 4-20% tris-glycine gel under non-reduced and reduced conditions. These two proteins were also treated with PNGase-F (NEB, MA) under the reduced condition to remove N-glycans and loaded on the gel to assess the impact of the glycans on the protein size. Gels were stained using Coomassie Blue and analyzed using a Bio-Rad G900 densitometer with Image Lab software.

### 2 Supplemental Figures

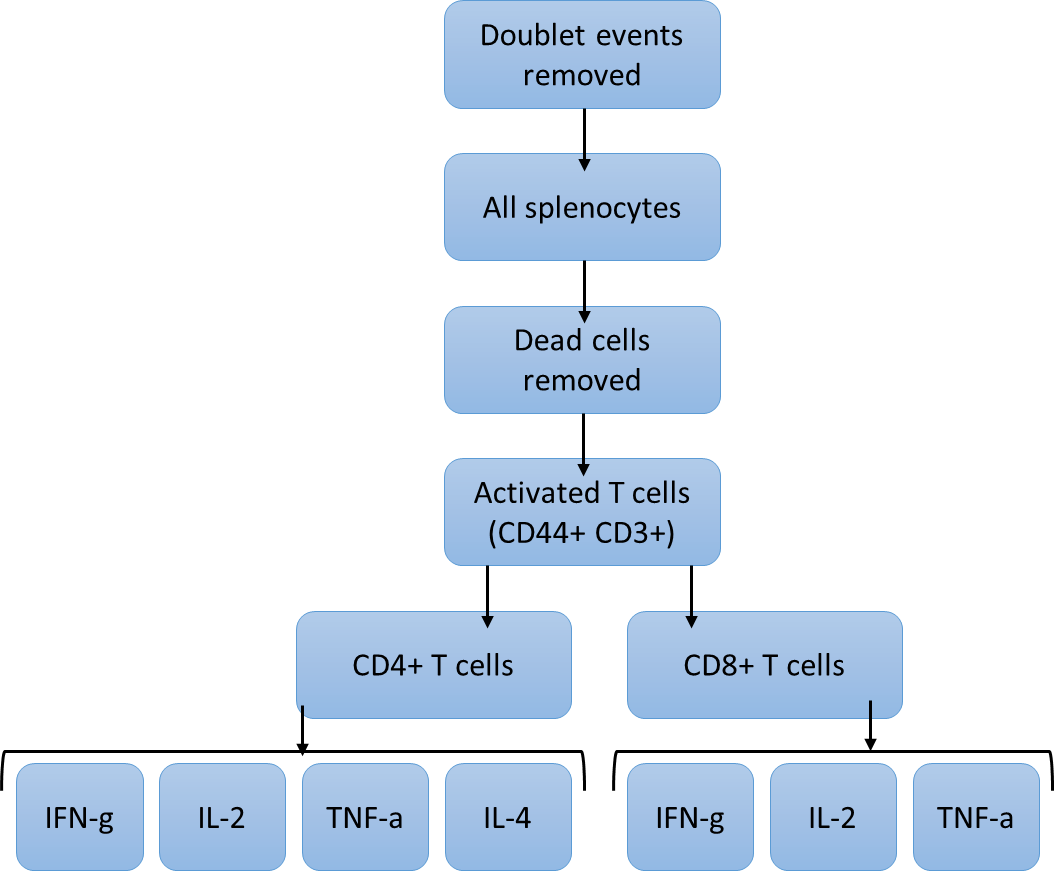

Supplemental Figure 1 : Gating strategy to investigate the cellular immune response in mice vaccinated with the SARS-CoV2-RBD219-N1-C1 vaccine

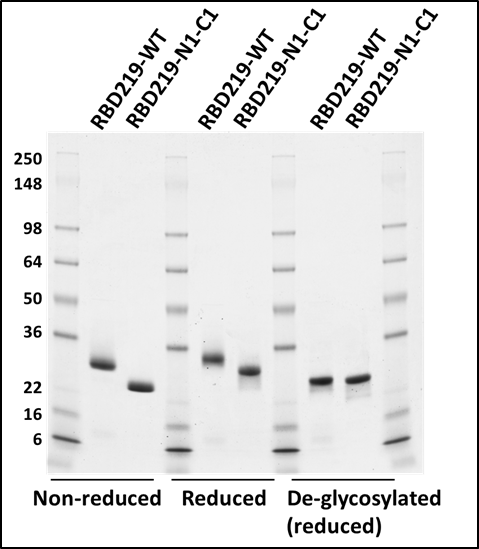

Supplemental Figure 2: Size evaluation of SARS-CoV-2 RBD219-WT and RBD219-N1-C1 using SDS-PAGE under non-reduced, reduced conditions, and after deglycosylation.

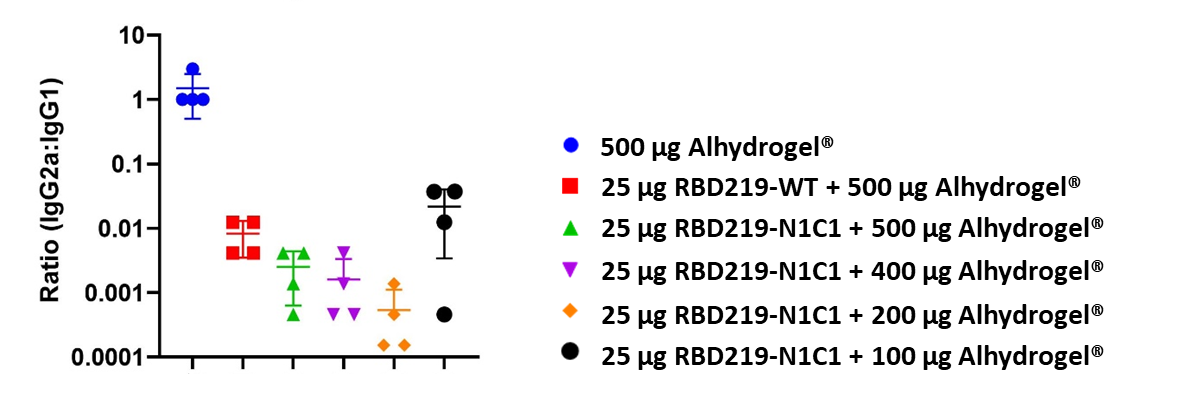

Supplemental Figure 3: Ratio of antigen-specific IgG2a:IgG1 in BALB/c mice (n =4) 43 days after the first immunization and 21 days after boost vaccination.
